# Liver fluke granulin promotes exosome-mediated crosstalk and cellular microenvironment conducive to cholangiocarcinoma

**DOI:** 10.1101/700427

**Authors:** Patpicha Arunsan, Apisit Chaidee, Christina J. Cochan, Victoria H. Mann, Toshihiko Tanno, Chutima Kumkhaek, Michael J. Smout, Shannon E. Karinshak, Rutchanee Rodpai, Javier Sotillo, Alex Loukas, Thewarach Laha, Paul J. Brindley, Wannaporn Ittiprasert

## Abstract

Crosstalk between malignant and neighboring cells contributes to tumor growth. In East Asia, infection with fish-borne liver flukes is a major risk factor for cholangiocarcinoma (CCA). The parasite secretes a growth factor termed liver fluke granulin (*Ov*-GRN-1), a homologue of the human progranulin (huPGRN), which contributes significantly to biliary tract fibrosis and morbidity during infection. Here, exosome-mediated transfer of mRNAs from the human cholangiocyte cell line (H69) was investigated following exposure to *Ov-*GRN-1, to naïve recipient H69 cells. To minimize the influence of endogenous huPGRN, the gene encoding huPGRN was inactivated using CRISPR/Cas9-based gene knock-out. Several huPGRN-depleted cell lines, termed ΔhuPGRN-H69, were established. These lines exhibited >80% reductions in levels of huPGRN transcripts and protein, both in gene-edited cells and within exosomes released by these cells. Profiles of exosomal RNAs (exRNA) from ΔhuPGRN-H69 for CCA-associated characteristics revealed a paucity of transcripts for estrogen- and Wnt-signaling pathways, peptidase inhibitors and tyrosine phosphatase related to cellular processes including oncogenic transformation. Several CCA-specific mRNAs including MAPK/AKT pathway members were induced by exposure to *Ov*-GRN-1. By comparison, estrogen, Wnt/PI3K and TGF signaling and other CCA pathway mRNAs were upregulated in wild type H69 exposed to *Ov*-GRN-1. Of these, CCA-associated exRNAs modified the CCA microenvironment in naïve recipient cells co-cultured with exosomes from ΔhuPGRN-H69 exposed to *Ov-*GRN-1, and induced translation of MAPK phosphorylation related-protein in naïve recipient cells comparing with control recipient cells. Exosome-mediated crosstalk in response to liver fluke granulin promoted a CCA-specific program through MAPK pathway which, in turn, established a CCA-conducive disposition.

## Introduction

Cholangiocarcinoma (CCA) describes malignancy arising from the biliary epithelium. CCA originates in the cholangiocyte, the specialized epithelial cell that lines the intrahepatic and extrahepatic bile ducts, except for the epithelial cells of the gallbladder. Many CCA are adenocarcinomas (1, 2). Although the causative agent for many cancers remains obscure including non-liver fluke infection-associated CCA, the principal risk factor in liver fluke-endemic regions is well established: infection with *Opisthorchis viverrini* and related parasites (3-6). Infection with *O*. *viverrini* is the principal risk factor for CCA in the Lower Mekong River Basin countries including Thailand, Lao PDR, Vietnam and Cambodia (5, 6). It has been estimated that 10% of people chronically infected with liver flukes will develop CCA (7). In regions endemic for opisthorchiasis, the prevalence of CCA can exceed 80 cases per 100,000 residents (8).

Helminth parasites communicate and interact at the host-parasite interface (9). Communication is facilitated by metabolic products secreted from the tegument and excretory tissues, including via exosomes (10, 11). The liver fluke *O*. *viverrini* releases proteins and other metabolites (12), which influence host cells including cholangiocytes in diverse ways (13-16). Whereas the full complement of metabolites released by this parasite remains generally to be investigated for roles of communication and disease, a secreted protein termed liver fluke granulin (*Ov-*GRN-1) has been the focus of increasing investigation. *Ov*-GRN-1 is a paralogue of human granulin, which like the human granulin stimulates cell proliferation and wound healing, and which appears to contribute to the pathogenesis of opisthorchiasis (17-21). We have exploited this link to more deeply explore the role of *Ov*-GRN-1 in pre-malignant lesions of the bile duct by undertaking programmed CRISPR/Cas9 knockout of the its cognate gene. Deep sequencing of amplicon libraries from genomic DNA of gene-edited parasites revealed Cas9-catalyzed mutations within the *Ov-grn-1* locus. Gene knockout rapidly depleted of transcription of *Ov-grn- 1* and depleted cellular levels of *Ov*-GRN-1. Furthermore, experimental infection of hamsters with the gene edited liver flukes resulted in markedly reduced disease even though gene-edited parasites colonized the biliary tract and developed into the adult developmental stage of the liver fluke. These findings confirmed a role for *Ov*-GRN-1 in virulence of the hepatobiliary morbidity characteristic of opisthorchiasis (22).

In the present report, we investigated exosome-mediated transfer of functional mRNAs to naïve recipient cells from cholangiocytes that had been exposed to liver fluke granulin. In response, exosome-mediated crosstalk promoted CCA-specific transcriptional profiles in the recipient cholangiocytes, including via MAPK signaling that, in turn, established a microenvironment supportive of carcinogenesis.

## Results

### Progranulin mutation in cholangiocytes by CRISPR knockout

The progranulin gene, *PGRN*, in the H69 cholangiocyte cell was targeted for CRISPR/Cas9-mediated programmed mutation. The programmed target site encodes the N-terminus and part of the granulin/epithelin module (GEM) of human PGRN (Fig. A, B). Wild type H69 cells were transduced with pLV-huPGRNx2 virions at > 5×10^*rOv*^ IFU/ml and, a day later, culture medium supplemented with puromycin at 300 ng/ml added to the virion-transduced cells with the goal of enrichment of transgenic cells exhibiting puromycin resistance (23). Three daughter cell lines, termed ΔhuPGRN-H69 lines 1, 2 and 3, exhibited > 70% reduction in levels of *huPGRN* transcripts and of protein as assessed by RT-PCR and western blot (WB), respectively (below). Thereafter, discrete amplicon libraries were constructed from each of the three daughter cell lines, from which 139,362, 107,683, and 179,122 sequence reads were obtained. The reads were analyzed using the CRISPResso pipeline to monitor programmed gene mutations and subsequent non-homologous end joining (NHEJ) mediated-repair and insertion-deletion (INDEL) profiles (24). This analysis revealed mutations in 32.3, 41.9 and 44.3% of the alleles in the lines, respectively. Similar INDEL profiles were seen among the three libraries: single nucleotide insertions, 6 and 2 bp deletions, and also other deletions from one to 10 bp in length were common at the position predicted for the programmed double stranded chromosomal break (Fig. 1, C-E, dashed line). Fig. 1E presents examples of the mutated alleles. This mutational landscape resembled profiles reported for chromosomal repair of CRISPR/Cas9 double stranded breaks by NHEJ in other mammalian cells (25). GenBank at NCBI assigned These sequence reads from amplicon-NGS libraries from the mutated *PGRN* locus of the ΔhuPGRN-H69 lines have been assigned the SRA accessions SRR9735031-34, BioProject ID PRJNA556235, and BioSample accession SAMN12344265.

**Figure 1.**
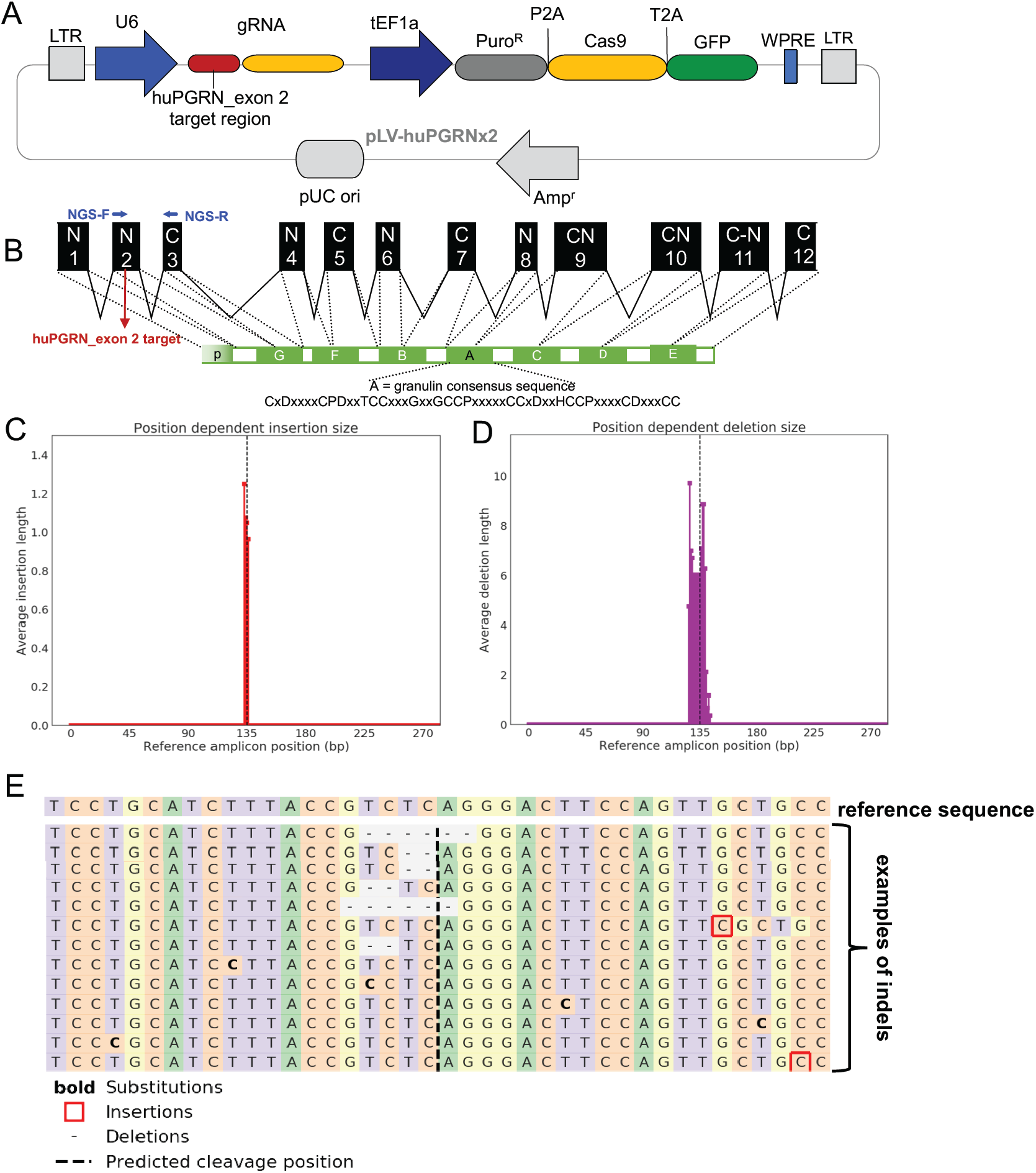
Programmed CRISPR/Cas9 mutation of the human progranulin gene. **Panel A**, Linear map of a pre-designed lentiviral CRISPR/Cas9 vector containing fused codon-optimized puromycin resistance marker (dark gray bar)-Cas9 (yellow bar) and green fluorescent protein (green bar) driven by the mammalian elongation factor alpha-1 promoter (dark blue arrow); guide RNA (gRNA) targeting human granulin exon 2 (red bar) is expressed from a single vector. The gRNA was driven by the human U6 promoter (blue arrow). The vector backbone includes the 5’- and 3’-long terminal repeats (LTR) of the HIV-1 provirus (light gray blocks). **Panel B**, Schematic representation of the partial human granulin gene precursor (huGRN) on chromosome 17: NC_000017.11 regions 44,345,086-44,353,106 (8,021bp) and protein structure. Nucleotide sequence in exon 2 encodes the N-terminus and part of the granulin/epithelin module (GEM) of progranulin; indicating locations of gRNA (4,417-4,438 nt; red colored-letter) predicted double-stranded break (DSB) (red arrow). **C-E**, On target INDEL mutations in ΔPGRN-H69 analysis by NGS libraries and CRISPResso bioinformatic platform. Frequency distribution of position-dependent insertions (red bars) (C) and deletion (magenta) (D) the major indels; 6 and 2 bp deletions at the programmed CRISPR/Cas9 cleavage site. Other minor mutations including 1 bp insertion (red square) or base substitution (bold) were observed further (>10 bp) predicted cleavage site (E).

### Growth and proliferation of PGRN knockout cell lines promoted by liver fluke granulin

Prolonged reduction in levels of huPGRN were observed for > 40 passages in three daughter ΔhuPGRN-H69 lines under puromycin treatment (not shown). By that point, differential huPGRN transcript fold change were stably reduced to ∼30% (29.8±4.7%) in ΔhuPGRN-H69 cells compared with levels in the parental, wild type cells (Fig. 2A) (*t*-test, *P*< 0.0001). When examined by WB, the levels of huPGRN in the ΔhuPGRN-H69 cells had fallen to 21.5±2.3% of those in the parent H69 cells, normalized against the human GAPDH reference (Fig. 2B) (*t*-test, *P*< 0.0001). A band at 64 kDa, the predicted mass of human granulin was evident in each of three biological replicates (Fig. 2B, lanes b1-3), as was the GAPDH signal at 36 kDa.

**Figure 2.**
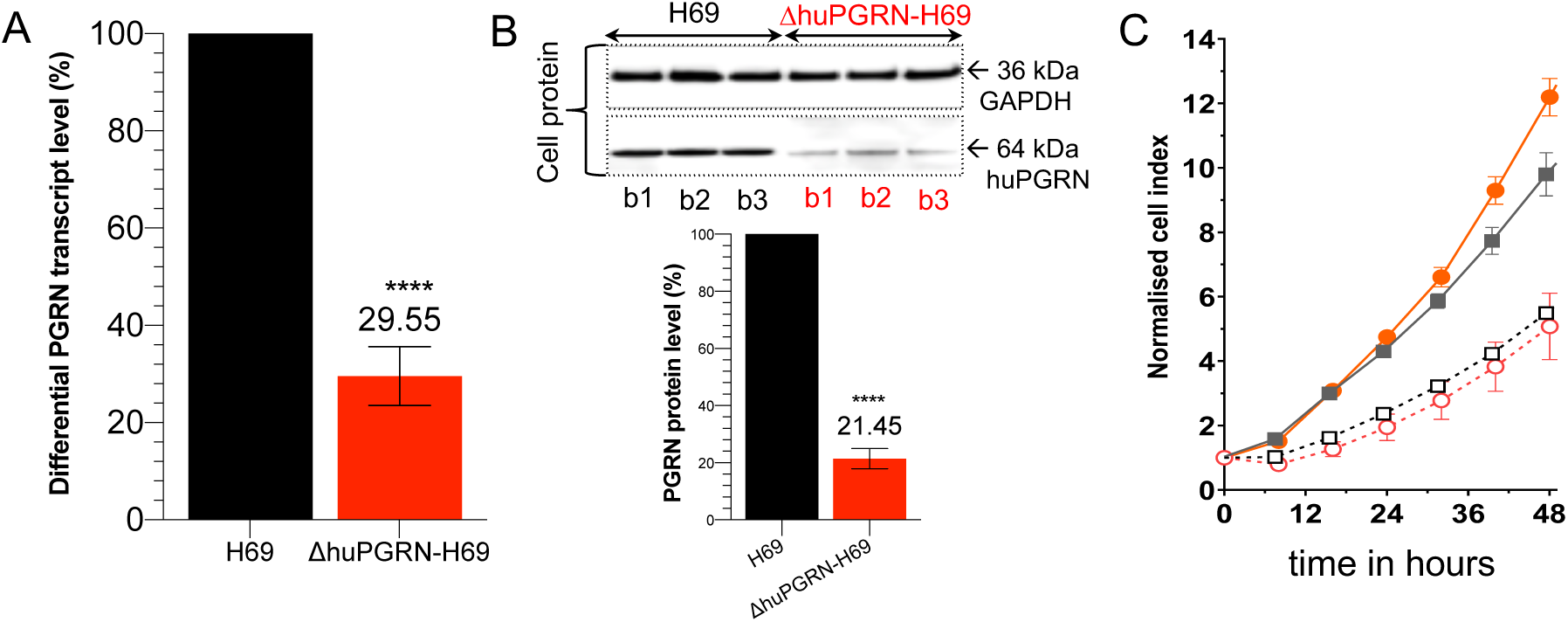
Reduction of huPGRN transcript and protein expression levels and cell proliferative effects of liver fluke granulin. **Panels A and B**. Reduction of human GRN transcription levels from ΔhuPGRN-H69 cell; red bar (∼70%) comparing with H69 reference (black bar). The huPGRN differential transcript after normalization with human GAPDH gene; mean ±SD, *n* = 3 (biological replicates); *P* < 0.0001 (****), unpaired *t*-test. Diminished level of huPGRN revealed by WB analysis using anti-PGRN antibody (64 kDa) compared with anti-GAPDH antibody (36 kDa). HuPGRN level in in ΔhuPGRN-H69 cells (red bars) was reduced by ∼80%. The GAPDH levels were stable within three biological replicates. These decreased levels of huPGRN were significantly different from H69 (black bars); *n* = 3 (biological replicates); *P* < 0.0001 (****), unpaired *t*-test. **C**. Real time monitoring of H69 vs ΔhuGRN-H69 cell proliferation before and after addition of liver fluke granulin. The lower normalized cell index (nCI) of ΔhuGRN-H69 cells (discontinuous red lines) compared with H69 cells (discontinuous gray lines) were monitored over 48 hours. The nCI of ΔhuGRN-H69 cells was recovered as in H69 cells nCI value after addition of liver fluke granulin at 100 nM for 24 h, and higher than H69 from 24 to 48 h. The nCI signals are shown as mean ± SD, for ≥3 independent experiments, and assessed using one-way ANOVA, *P* < 0.001.

To assess the consequence of loss of huPGRN following the programmed gene knockout, cellular proliferation and the mitogenic activity of recombinant *Ov*-GRN-1 (r*Ov*-GRN-1) was monitored using the xCELLigence approach. This label-free cell-based assay system uses culture plates bearing microelectrodes for non-invasive measurement during cell culture (26). The rate of H69 and ΔhuPGRN cell growth was monitored for 48 hours and determined by calculating the slope of the line between readings, which were collected at 20 min intervals during the assay. Both the H69 and the daughter mutant ΔhuPGRN grew and proliferated at a similar rate for 48 h, indicating that the programmed knockout of *huPGRN* was not detrimental to normal cell growth. Next, the wild type and mutant H69 cell lines were pulsed with r*Ov*-GRN-1 at 100 nM. Upon addition of liver fluke granulin, the rates of proliferation of ΔhuPGRN-H69 and H69 were similar over the first 24 h, after which ΔhuPGRN-H69 cells grew significantly quicker than H69 until 48 h (Fig 2C, solid lines) Liver fluke granulin induced significantly faster growth in both H69 and ΔhuPGRN compared to control groups of both cell lines not exposed to r*Ov*-GRN-1, as described previously for H69 and other cells (one-way ANOVA; *P* < 0.001) (17, 21). Figure 2C presents the mean (± SD) slope of three experiments performed at seven intervals from 0 to 48 h, for each of the parental H69 lines cells (gray) and the daughter ΔhuPGRN lines (red).

### H69 cholangiocytes released exosomes

Exosome particles recovered from culture supernatants of H69 cells investigated. Dot-blot based biochemical characterization (Exo-Check Array) revealed the following exosomal markers with intensity (gray scale) between 10-70: flotillin-1 (FLOT-1), intercellular adhesion molecule 1 (ICAM), ALG-2-interacting protein 1 (ALIX), CD81, epithelial cell adhesion molecule (EpCAM), annexin V (ANXAS) and tumor susceptibility gene 101 (TSG101), while the positive control showed intensity >100. The purified exosomes were negative for *cis*-Golgi matrix protein (GM130) with absence of intensity as negative control, indicating the absence of contaminating cellular debris (Fig. 3B). WB analysis of the exosomes confirmed the expression of CD9 and CD81, hallmark surface markers of exosomes (26, 27), with chemiluminescent signals on the blot at 24 kDa and 26 kDa, respectively (Fig. 3C). Confocal microscopical observation revealed exosome particles that were probed with anti-CD81 labeled with fluorophore 488 (green) surrounding DAPI-stained nuclei (blue) of H69 cells (Fig. 3A). Exosome particle size distribution ranged from 55 to 80 nm (not shown). These findings confirmed the identity of the particles released from the H69 cells as exosomes.

**Figure 3.**
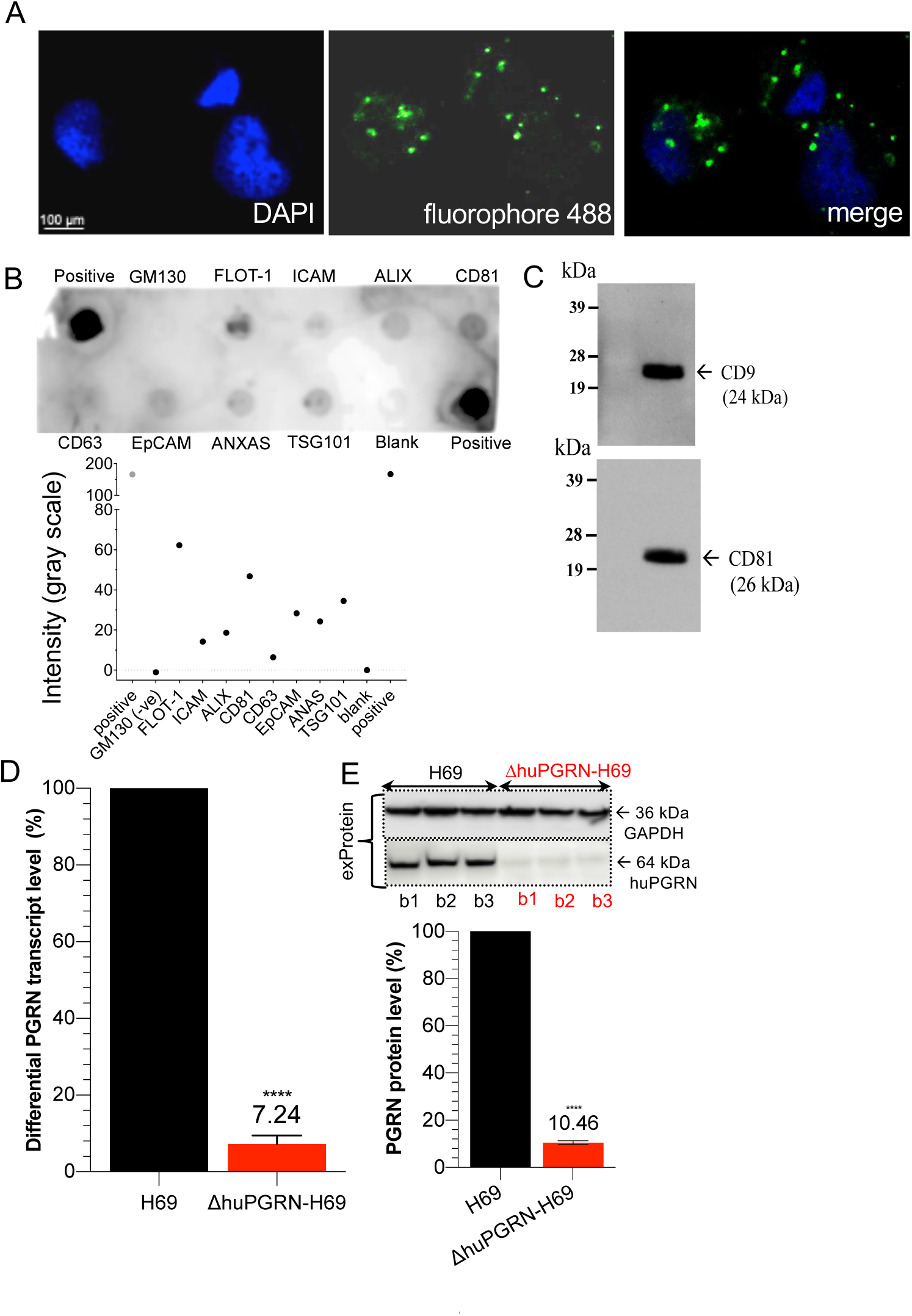
Exosome derived H69 cell characterization and exosomal PGRN expression. **Panel A**, the visualization of exosomes from H69 cells by *in situ* hybridization using fluorophore 488-labeled anti-CD81 antibody (green dot, panel A) around DAPI-stained cell nuclei (blue color). The H69-derived exosome particle sizes ranged from 55 to 80 nm. The exosomal protein composition were determined using the Exosome Antibody Array (SBI System Biosciences). **B**, dark spots indicate presence of the target protein (upper panel). Absence of a spot for GM130 indicated absence of cellular contaminants. The intensity in gray scale was also plotted (lower panel). **C**, WB analysis for the exosome-specific markers; strongly positive signal for CD9 and CD81 were found. **D, E**, both the exRNA and exProtein from ΔhuPGRN-H69 showed reduction of ∼90% differential transcript levels [red bar, panel D] after normalization with GAPDH in comparison to H69 [black bar] (red bar, panel D). The levels of huPGRN were significantly reduced (∼90%); unpaired *t*-test, *P* < 0.0001 (****), *n* = 3.

### Programmed knockout of PGRN depleted exosomal levels of granulin

The programmed mutation induced INDELs at exon 2 of the huPGRN locus negatively impacted translation of the huPGRN within the protein complement of the exosomes. Specifically, we determined the expression of huPGRN exosomal mRNA (exRNA) and protein (exProtein) released by each of three replicate lines of puromycin resistant cells, at passage number 20 in each case. By this point, exosomes from ΔhuPGRN-H69 cells included 7.2±2.6% only of the levels of *huPGRN* transcripts and 10.5±2.0% huPGRN compared to exosomes from H69 cells, after normalization with GAPDH (Fig. 3D, E) (unpaired *t*-test, *P* < 0.0001, both for transcription and protein expression). Expression of huPGRN was revealed by probing the WB with an anti-huPGRN antibody (target mass, 64 kDa) compared with anti-GAPDH antibody (36 kDa). Levels of GAPDH were similar both in cell lysates and exosomal proteins of the parental H69 and daughter ΔhuPGRN cells (Fig. 3E).

### CCA-related profiles of transcripts in exosomes shed liver fluke granulin-exposed cholangiocytes

To investigate if liver fluke granulin stimulated H69 cells secrete exosomes containing CCA-related transcripts, the huPGRN gene was mutated by CRISPR/Cas9 knockout (above). As noted, this gene knockout was undertaken in order to minimize background function of endogenous progranulin. The exosome-exposed ΔhuPGRN-H69 cells showed similar CCA-related mRNA base line profiles as seen with the CCA-array panel of 88 targets (Bio-Rad) probed with the H69 exRNAs. However, 10 exRNAs were absent from exosomes shed from ΔhuPGRN-H69 cells following gene knockout; specifically, dihydropyrimidine dehydrogenase (DPYD), estrogen receptors (ESR) 1 and 2, lysine-specific demethylase 3A (KDM3A), lymphoid enhancer-binding factor-1 (LEF1), protein tyrosine phosphatase (PRKC9), non-receptor type 13 (PRIN13, a Fas-associated phosphatase), serpin peptidase inhibitor, clade A, member 3 (SERRINA3), serpin peptidase inhibitor, clade E, member 2 (SERRPINE2), and SRY-related HMG-box 11 (SOX11) (Fig. 4A). By contrast, these all of these transcripts were present in exRNA of H69 cells. DPYD, ESR1, ESR2, KDM3A and SOX11genes participate in pyrimidine catabolism, hormone binding, DNA binding and activation of transcriptions and activate transcription factor, respectively. LEF1encodes a transcription factor that is involved in Wnt signaling. PTPN13 is a member of protein tyrosine phosphatase family, which regulates a variety of cellular processes including oncogenic transformation (28). The PRINA3 and SERPINA3 genes encode inhibitors of proteases and peptidases.

**Figure 4.**
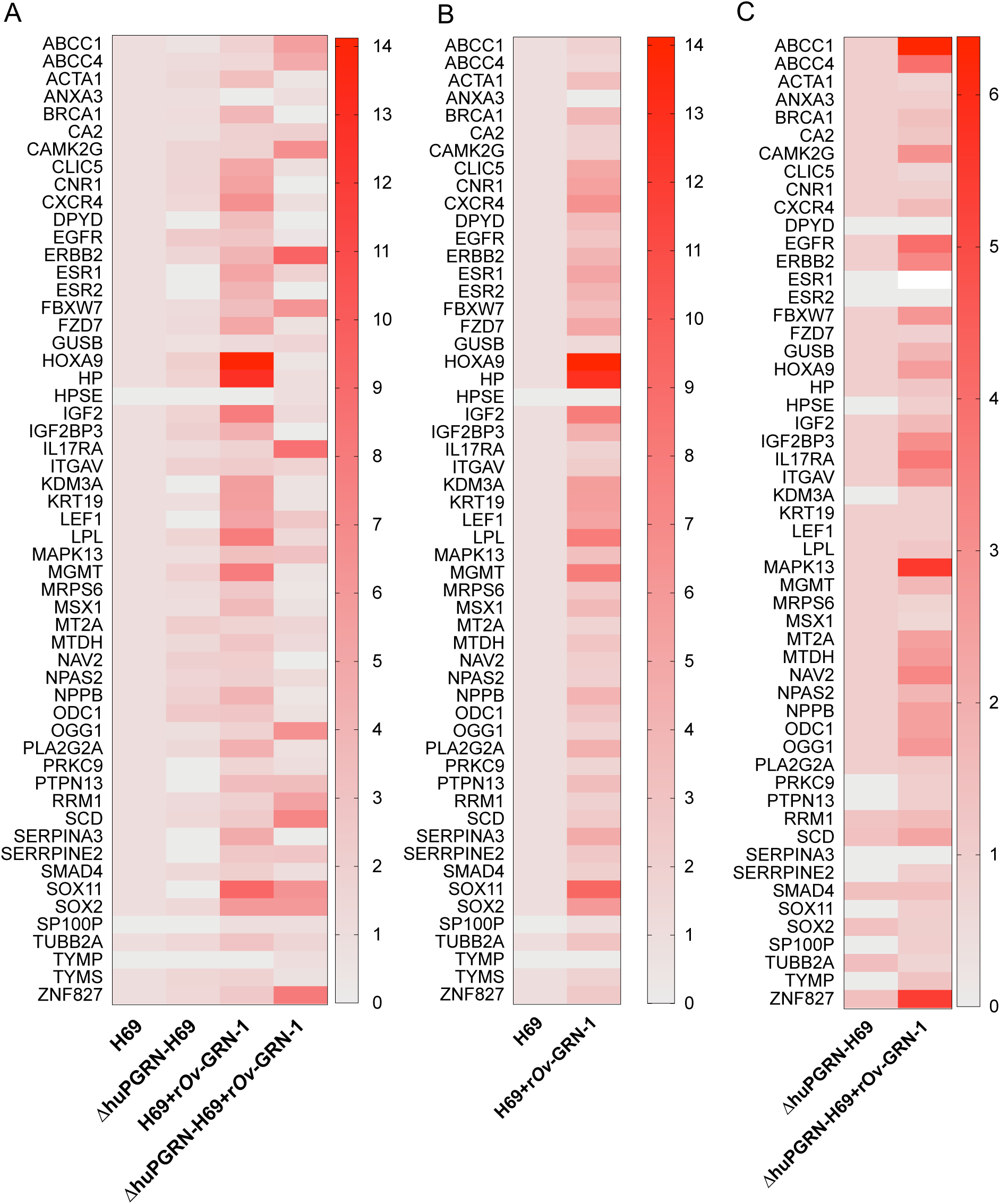
Induction of CCA-related mRNAs carrying cholangiocyte derived-exosome particles after exposure to liver fluke granulin. **Panel A**. The heat map plotted from the differential transcript fold change by PrimePCR^™^ CCA-related gene Array (Bio-Rad) using H69 as the reference from 1) ΔhuPGRN-H69 *vs*. H69 2) H69 with r*Ov*-GRN-1 treatment and 3) ΔhuPGRN-with r*Ov*-GRN-1 treatment. Similar quantities of exRNAs from each group were probed individually with 88 CCA targets in triplicate; there were slightly different (* *P* = 0.0629 by two way ANOVA gene profiles (55 from 88 transcripts were detected) in ΔhuPGRN-H69 vs H69 (panel A) by absent of DPYD, ESR, KDM3A, LEF1, PRKC9, PTPN13, SERPINA3, SERRPINE2 and SOX11. However, most of these genes were recovered after exogenous GRN treatment of ΔhuPGRN-H69 (panels A, C). The CCA mRNA profiles of exosomes derived from ΔhuPGRN-H69 and H69 were markedly dissimilar following exposure to liver fluke granulin with *** and **** *P* = <0.0001, 2way ANOVA (panels A and B). Panel C, shown the CCA-related exRNAs after exposure to liver fluke granulin in ΔhuPGRN-H69 comparing with its mRNA profile (**** *P =* 0.0001 with two-tailed, paired t test). The statistical significance was calculated from differential fold changes in transcription as per the manufacturer’s software following by Prism v8. The liver fluke granulin (r*Ov*-GRN-1) treatment group of cells showed 1-15 folds level of induction of transcription. The ‘1’ indicates baseline level expression, and ‘0’ indicated the absence of expression. The heat map was plotted from the average transcript level (*n* = 3) using Prism v8.

The profiles of exRNA (from three pooled biological replicates) were examined. Following exposure of ΔhuPGRN-H69 to liver fluke granulin, seven transcripts were detected based on the array panel, specifically ESR1, KDM3A, LEF1, PRKC9, PTPN13, SERRPINE2, and SOX11 (Fig. 4A, normalized with H69; Fig. 4C, normalized with ΔhuPGRN-H69).

### Specific CCA pathways impacted by liver fluke granulin-induced activation

There were 52 and 43 from 88 CCA-related genes expressed from H69 and ΔhuPGRN-H69 derived-exosomes following treatment with r*Ov*-GRN-1, respectively. The transcript profiles H69 and ΔhuPGRN-H69 derived-exosomes were dissimilar (Fig. 4). For exRNA from H69 cells, elevated levels were induced (4.1-to 14.1-fold changes) for genes encoding the chloride intercellular channel 5 (CLIC5), cannabinoid receptor (CNR1), chemokine (C-X-C-motif) receptor 4 (CXCR4), epidermal growth factor receptor (EGF) family (ERBB2), estrogen pathway (ESR1, ESR2), Frizzled 2-Wnt receptor (FZD7), homeobox A9 (HOXA9), haptoglobin (HP), insulin-like growth factor (IGF), insulin-link growth factor 2 mRNA binding protein 3 (IGF2BP3), zinc finger protein (KDM3A), keratin family (KRT19), transcription factor involved in the Wnt signaling pathway (LEF1), lipoprotein lipase (LPL), MAP kinase family (MAPK13), *O*^6^-methylguanine-DNA-methyltransferase (MGMT), natriuretic peptide B (NPPB), phospholipase A2 (PLA2G2A), peptidase inhibitor (SERPINA3), SRY-related HMG-box (SOX 2 and SOX11) family involves in the regulation of embryonic development and in the determination of cell fate (Fig. 4A, B). By comparison, r*Ov*-GRN-1 activated-ΔhuPGRN-H69 (i.e. without endogenous progranulin) showed elevated levels of transcripts (4.8-to 9.5-fold change) encoding ATP-binding cassette transporter (ABCC1, ABCC4), serine/threonine protein kinase family (CAMK2G), ERBB2 enhancing kinase-mediated activation, F-box protein family (FBXW7), proinflammatory cytokine 17A (IL17A), reductase subunit M1 (RRM1), MAPK13, enzyme responsible for the excision of 8-oxoguanine, enzymes involved in fatty acid biosynthesis (SCD), protein tyrosine phosphatase family involved in cell growth differentiation mitotic cycle and oncogenic transformation (PTPN13), SOX2, SOX1, and zinc finger protein ZNF827 involved in telomere chromosome remodeling (Fig. 4A, C). Based on these findings, at least two exosome transcripts, ERBB2 and MAPK13 were identified as participants of the MAPK signaling pathway and detected with high transcript fold change (>4) in exosomes derived from ΔhuPGRN-H69 cells following activation by liver fluke granulin. These findings concur with a central role hypothesized for the MAPK pathway in liver fluke infection induced-cholangiocarcinogenesis (21, 29).

### Human progranulin supplements impact of liver fluke granulin on cholangiocarcinogenesis

Analysis using the CCA PrimePCR Array (Bio-Rad) revealed that 41 of the 55 CCA-related genes induced transcription by two-to four-fold. These 41 genes included actin 1 (ACTA1), breast cancer 1 (BRCA1), epidermal growth factor receptor (EGFR), F-box and WD40 repeat domain containing-7 (FBXW7), MAPK13, mitochondrial ribosomal protein S6 (MRPS6), msh home box 1 (MSX1), metadherin (MTDH), neuron navigator 2 (NAV2), transcription factor (NPAS2), ornithine decarboxylase 1 (ODC1), RRM1, SCD, SERRPINE, TGF-beta signaling family member (SMAD4), tubulin β2α (TUBB2A), ZNF827. Additional genes with elevated levels of transcripts in exRNA (> 4-fold change) were noted above. Based on these findings, the endogenous growth factor progranulin appeared to supplement the action of liver fluke granulin in the cholangiocyte in establishing a carcinogenesis-conducive transcriptional profile. This outcome may mimic events during natural infection with *O*. *viverrini* where the liver fluke residues within the bile duct, secretes liver fluke granulin which is, in turn, taken up by cholangiocytes (17) and, which contributes to biliary tract inflammation and fibrosis, and ultimately to cholangiocarcinoma (6, 21, 30).

### Exosomes secreted from liver fluke granulin activated donor cell transfer CCA-MAPK-related functional mRNAs to naïve recipient cells

To investigate paracrine communication involving transfer of exosomal mRNA from donor liver fluke granulin treated-ΔhuPGRN-H69 cells to naïve neighboring cells (H69 or -ΔhuPGRN-H69 cells), recipient cells were co-cultured with exosomes collected from r*Ov*-GRN-1-activated ΔhuPGRN-H69 (exosome^*rOv*^). Extracellular exosomes were endocytosed by other cells, as revealed by PKH-26 labelling (31), and delivered their cargo to the recipient cells (Fig. 5A, 5B). Treatment with Pitstop 2 (32) blocked endocytosis of exosomes by H69 (Fig. 5C). H69 cells initiated endocytosis of exosomes within 30 min, and endocytosed ∼90% of cell population within 90 min (Fig. 4). The cargo of exosome^*rOv*^ CCA-related mRNAs was functional and stimulated changes in gene expression in naïve recipient cholangiocytes. Focusing on MAPK pathway phosphorylation from among these changes in gene expression, we employed a commercial human MAPK phosphorylation array that included 17 kinase phosphorylation sites. Following exosome^*rOv*^ co-culture, recipient cells were collected and processed. Seven proteins - GSK3a, MKK3, mTOR, p38, p53, P70S6K and RSK1 - were increased significantly by exosome^*rOv*^ treated-ΔhuPGRN-H69 compared with the control group (Fig. 5D -F) (paired *t* test, *** *P =* 0.0009, t = 4.050, df=16, number of pairs = 17). The pattern of protein induction was similar in exosome^*rOv*^-treated H69, except for GSK3a (not significant) (Fig. 5D) (paired *t* test, * *P =* 0.0464, *t* = 2.159, df=16, n = 17). The indirect amount of each protein was measured with normalized signal intensity (RFU); more of these seven proteins were present in H69 than ΔhuPGRN-H69 cells. For example, MKK3 showed ∼ 4,000 RFU and ∼3,700 RFU in H69 and ΔhuPGRN-H69 control groups, respectively. However, the activation of MAPK phosphorylation by mRNAs from exosome^*rOv*^ exhibited a similar protein profile. These findings confirmed paracrine signaling among cholangiocytes, which was accomplished by exosome-mediated message transfer from liver fluke granulin-exposed cells to naïve cholangiocytes.

**Figure 5.**
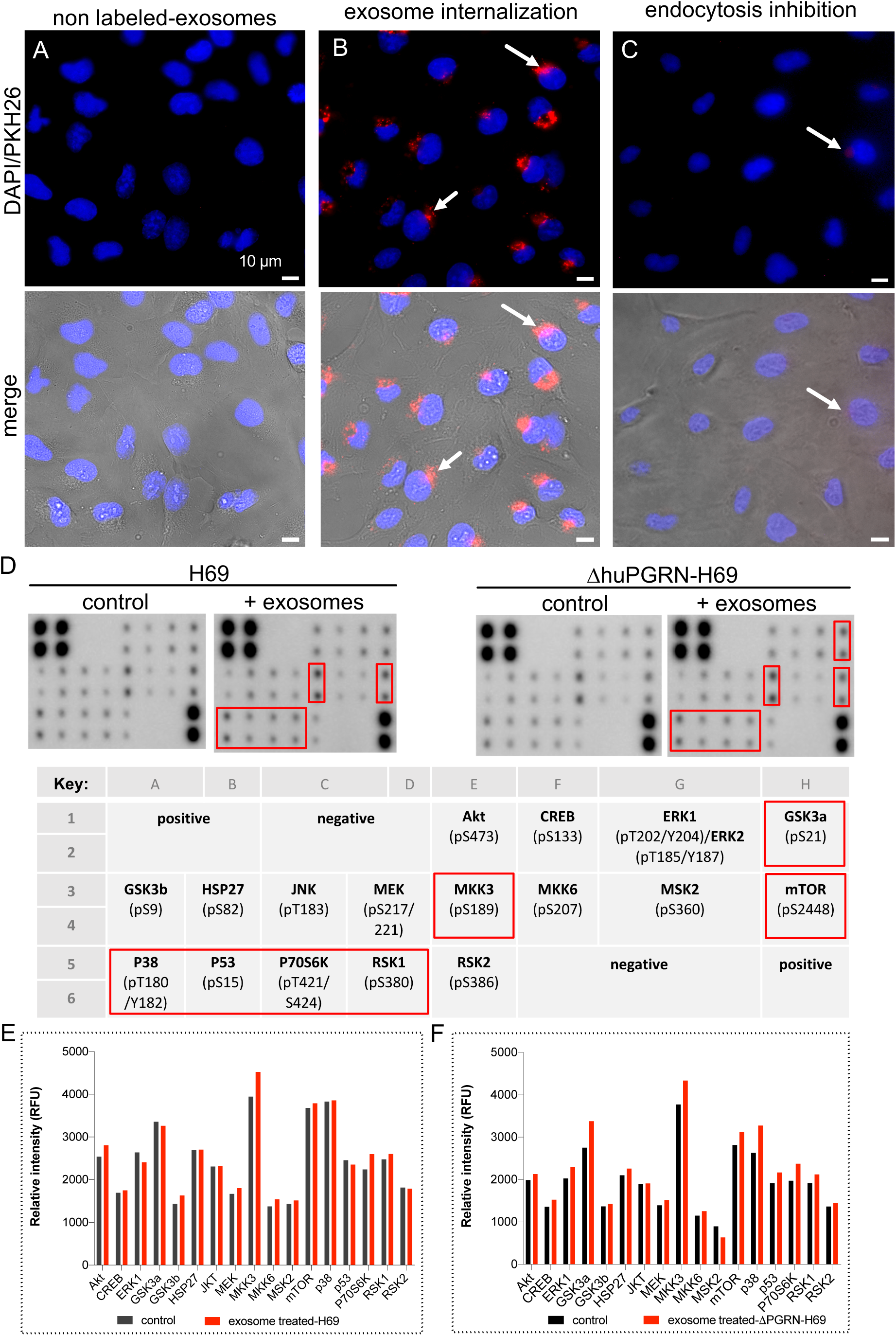
Uptake of extracellular exosomes and paracrine transfer of functional mRNAs from donor to naïve recipient cells. **Panels A, B**, representative fluorescence micrographs showing uptake of PKH26-labeled exosomes into recipient cholangiocytes; blue—nuclei staining with the NucBlu Live Cell Stain ReadyProbe, red—PKH26-labeled exosomes. Pitstop 2 blocked clathrin-dependent endocytosis of exosomes (**C**). Merged images shown in lower panel with bright field. Magnification, 40×; scale bar, 10 µm. **D, E**, cholangiocytes were grown to 80% confluence after which they were co-cultured with exoxomes^*rOv*^ for 24 h. The cell lysate (500 ng) was used for 17 targets on the human MAPK phosphorylation antibody array (Abcam), and signal intensity for each spot quantified by densitometry (**D**). Red boxes represent the predominant proteins that significantly higher than control groups namely, GSK3a, MKK3, mTOR, p38, p53, p70S6K and RSK1. The MAPK phosphorylation protein pattern revealed statistically significant differences in comparison with its control (*P* < 0.05, paired *t*-test)). Data represent the average of two individual sets per sample.

## Discussion

In East Asia, infection with fish-borne liver flukes is a major risk factor for cholangiocarcinoma (CCA). The liver fluke *O*. *viverrini* secretes a growth factor termed liver fluke granulin, *Ov*-GRN-1. This growth factor is a paralogue of progranulin (PGRN), which is a secreted, cysteine-rich glycoprotein that regulates cell division, survival, motility and migration. Progranulin participates in embryonic development, wound repair, and cancer. Mutations in the human progranulin (huPGRN) gene are associated with a spectrum on neurological disorders (33).

The liver fluke paralogue, *Ov*-GRN-1, stimulates cell proliferation, wound healing, and angiogenesis *in vitro* and contributes to the pathogenesis of opisthorchiasis *in vivo* (17-22). We have exploited this link infection to explore the role of the secreted liver fluke granulin in pre-malignant lesions by undertaking programmed CRISPR/Cas9 knockout of the *Ov*-GRN-1 gene from the genome of *O*. *viverrini*. Infection with *Ov*-GRN-1 knockout worms results in markedly reduced disease, which confirmed the key role for this liver fluke granulin in hepatobiliary morbidity during opisthorchiasis (22).

Crosstalk between malignant and neighboring cells contributes to tumor growth (34-37). To investigate crosstalk among cholangiocytes during liver fluke infection, here we focused on exosome-mediated transfer of mRNAs from cultured cholangiocytes following exposure to liver fluke granulin. To minimize the influence of endogenous huPGRN, the gene encoding huPGRN was inactivated in H69 cells using CRISPR/Cas9-based gene knock-out. Cholangiocytes of the resulting mutant ΔhuPGRN-H69 lines grew more slowly than the parental H69 cells. Notably, however, the cell growth recovered and increased following exposure to liver fluke granulin, and indeed liver fluke granulin-treated ΔhuPGRN-H69 cells grew faster than H69 between by 24 and 48 hours later.

Analysis using a CCA gene array indicated the cooperation of CCA genes with the exosomes after exposure to liver fluke granulin, in both the H69 and ΔhuPGRN-H69 cells, highlighting the importance of signaling pathways related to intracellular communication. The exRNA expression pattern between H69 cells activated by exposure to liver fluke granulin and, functioning together with human progranulin, induced CCA mRNA induction (41 from 55 genes). By comparison, the exRNA pattern from ΔhuPGRN-H69 (where levels of endogenous huPGRN was markedly diminished) was substantially different (17 from 55 genes). This finding supports a role for liver fluke granulin during opisthorchiasis-induced carcinogenesis, involving activation of the MAPK pathway (17, 38). Moreover, the MAPK exRNAs were transferred by endocytosis to, and were functional in, naïve cholangiocytes. Endocytosis of exosomes by the H69 and ΔhuPGRN-H69 recipient cells was accomplished within 90 min. Subsequent *in vivo* translation of MAPK exRNA from exosomes^*rOV*^ was confirmed for GSK3a, MKK3, mTOR, p38, p53, p70S6K and RSK. In overview, exosome-mediated crosstalk in response to liver fluke granulin promoted CCA-specific programs including via MAPK signaling that, in turn, may contribute to the CCA-conducive microenvironment.

Cholangiocytes express IL-6 and IL-8 through the TLR4-NF-kB and MAPK signaling pathways (39). The MAPK pathway is central to cholangiocarcinogenesis activated by *Ov*-GRN-1 during *O*. *viverrini* infection (17, 38). For CCA development, oncogenic signaling pathways (40) participate in many steps of carcinogenesis (41-43). Wnt/β-catenin signaling stimulates the pathobiology associated with CCA (44), with Wnt/β-catenin signaling involved in the inflammation-associated CCA. Suppression of Wnt/β-catenin pathway signaling negatively impacts development of CCA (44). Moreover, apoptosis, growth, cellular intercommunication, and angiogenesis of CCA cells related with receptor tyrosine kinase (RTK) signaling that revealed the multiple kinases associated with PI3K/AKT, Wnt/β-catenin, MAPK, and/or JAK/STAT signaling pathways (45-47). In addition, GRN development affects the activation of the MAPK signaling pathway (48), highlighted by its upregulated of PGRN during hepatocellular carcinoma and as the gene target of microRNA-140-3p (miR-140-3p). Overexpression of miR-140-3p disrupts the stimulation in the MAPK signaling pathway by inhibition of PGRN expression, leading to phosphorylation of ERK and p38. Indeed, in this situation, c-Jun N-terminal kinase (JNK) is suppressed which, in turn, inhibits migration and invasion by tumor cells (49).

Endogenous progranulin supplemented the action of liver fluke granulin in establishing a carcinogenesis-conducive transcriptional profile. This outcome may mimic events during natural infection with *O*. *viverrini* where the liver fluke residues within the bile duct, secreting liver fluke granulin which, in turn, is taken up by cholangiocytes (17) and which contributes to biliary tract inflammation and fibrosis, and ultimately to cholangiocarcinoma (6, 21, 30). Here, transcription levels of the MAK13 and SOX2 were up-regulated following exposure of naïve wild type or the ΔhuPGRN cells to exosomal RNAs from ΔhuPGRN-H69 cells that had been exposed to liver fluke granulin. By contrast, SOX11 was up-regulated in the wild type cells. Perturbation in the MAPK and Wnt signaling pathways that are activated in these cells may reflect the natural history of development of CCA (44) where SOX2 expression is upregulated in the precursor cell and supports self-renewal of the transformed cell (50-52). SOX11 plays a role in the tumor cell progression and in maintenance of the protein complex associated with the Wnt and other signaling pathways that mediate angiogenesis (53-55).

Infection with *O*. *viverrini* appears to induce global epigenetic deregulation in the presence of chronic inflammation, provoked by the mechanical damage inflicted by the feeding and by other activities of the parasite, including secreted factors such as liver fluke granulin, thioredoxin peroxidase and, as shown here, extracellular vesicles that modulate the gene expression profiles (12, 16, 21, 22). We established a line of the H69 cholangiocyte with markedly diminished levels of endogenous progranulin, using CRISPR/Cas9-mediated programmed gene knockout. The findings presented here with the mutant H69 cell confirmed that exosome-mediated cellular communication, induced by exposure of cholangiocytes to liver fluke granulin, modulated conserved signaling pathways in ways reflective of cholangiocarcinogenesis. Cholangiocarcinogenesis induced by chronic opisthorchiasis and continuous exposure to liver fluke granulin may act through a RTK signaling, perhaps signaling via ephrin pathway receptor (56), and the interconnections of the MAPK and Wnt/β-catenin pathways.

## Materials and Methods

### Cell lines

The human nonmalignant immortalized cholangiocyte cell line (H69) was cultured in H69 complete medium; Ham’s F12 nutrient mixture (25 µg/ml adenine, 5 µg/ml insulin, 1 µg/ml epinephrine, 0.62 µg/ml hydrocortisone, 1.36 µg/ml T3-T, and 10 ng/ml epidermal growth factor (EGF), Dulbecco’s Modified Eagle Medium [DMEM] (Gibco), DMEM/F-12 (Sigma-Aldrich, St. Louis, MO) media containing 10% fetal bovine serum (FBS) and 1*×* penicillin/streptomycin (pen/strep), as described (21, 57, 58). H69 cells was maintained in humidified incubator in 5% CO_2_ in air at 37°C. H69 cells were *Mycoplasma*-free as established using the Lookout Mycoplasma PCR detection kit (Sigma-Aldrich) and authenticated using short-tandem repeat profiling (ATCC, Manassas, VA).

### Lines of progranulin knockout cholangiocytes

To mutate and disrupt the human progranulin gene, *huPGRN*, with the aim of minimizing or eliminating endogenous granulin in H69 cells that might obfuscate interpretation of the effect of liver fluke granulin, we employed a pre-designed lentiviral CRISPR/Cas9 approach - ‘All in One CRISPR/Cas9 vector system (Sigma-Aldrich). Our modified vector encoded a guide RNA targeting human PGRN (huPGRN) exon 2, which encodes granulin-epithelin precursor (GEP); 5’-cctgcaatctttaccgtctc-3’ of on chromosome 17: NC_000017.11 regions 44,345,086-44-353,106 driven by U6 promoter, and the elongation factor 1*α* promoter to drive the expression of fusion proteins of the puromycin *N*-acetyl transferase from *Streptomyces alboniger* (puromycin resistance marker, Puro^R^) (23), Cas9 endonuclease from *Streptococcus pyogenes*, and green fluorescent protein (GFP), in that order. The cargo genes of the vector are flanked by the long tandem repeat (LTRs) of the lentivirus (Fig. 1A). Competent *E*. *coli* cells were transformed with the plasmid vector, termed pLV-huPGRNx2, and maintained in LB broth, 100 µg/ml ampicillin.

Virions were derived following transfection of human 293T cells producer cells, using FUGENE HD transfection reagent (Promega, Madison, WI), with pLV-huPGRNx2 construct and the MISSION^™^ lentiviral packaging kit (Sigma-Aldrich), as described (59). Pooled culture supernatants containing pseudotyped virions were collected at 48 to 72 h after transfection of 293T cells, clarified by centrifugation at 500×*g* for 10 min, and passed through a Millipore 0.45 µm pore membrane (Steriflip-GP, Millipore). Virions were concentrated using Lenti-X concentrator (Takara Bio USA, Inc., Mountain View, CA) after which titers ere measured by Lenti-X-GoStix Plus (Takara Bio). For programmed knock-out of *huPGRN* gene, ∼350,000 H69 cells were exposed to 500 µl of pLV-huPGRNx2 virion (∼5×10^5^ infectious units [IFU]/ml) in 2.5 ml complete H69 medium in 6-well plates. One day later, the medium was replaced with medium supplemented with puromycin at 300 ng/ml for the selection and enrichment of cells carrying the proviral form of the gene-editing virus. (Survival of H69 cells in puromycin ranging from 50 to 400 ng/ml was tested at the outset, aiming to define a concentration to inhibit survival of these cholangiocytes. H69 cells were killed by puromycin within 48 hours in at 300 ng/ml puromycin [not shown].) Gene edited cells were maintained in parallel with H69 cells for 72 h, by which point the control H69 cells had died. About 5 -10% of cells survived and entered clonal amplification. Surviving cells were cultured in complete H69 medium supplemented with 300 ng/ml puromycin at 300 ng/ml for 20 passages before genotyping. Discrete biological triplicates were undertaken to establish three puromycin-resistant *huPGRN* knock-out cell lines (b1, b2 and b3 in Fig. 2), which exhibited >70% reductions in *huPGRN* transcript and protein levels (Fig. 2A, B). The lines are termed ΔhuPGRN-H69 lines.

### Programmed mutation

Following lentiviral transduction and maintenance in 300 ng/ml puromycin, surviving cells were expanded through successive passages. After passage 20, genomic DNAs were extracted from pools of the enriched, puromycin resistant ΔhuPGRN-H69 cells using DNAzol (Molecular Research Center). We amplified the targeted region of exon 2, huPGRN using the NGS primer pair; forward primer 5’-GACAAATGGCCCACAACACT-3’ and reverse primer 5’-GCATAAATGCAGACCTAAGCCC-3’ (Fig 1B) flanking expected double strand break (DSB) by CRISPR/Cas9 system. The DNA samples were processed for on-target insertion-deletion (indel) investigation by next generation sequencing (NGS), using the Ion Torrent Personal Genome Machine (ThermoFisher). The sequencing libraries were prepared from 10 ng DNA using the Ion Torrent Ampliseq kit 2.0-96 LV (ThermoFisher), following the manufacturer’s instructions. The DNA was bar-coded using the Ion Xpress Barcode Adapters kit and quantified by quantitative PCR using the Ion Library TaqMan Quantitation Kit (ThermoFisher) after purification of libraries by Agencourt AMPure XP beads (Beckman). Emulsion PCR was performed using the Ion PGM Hi-Q View OT2 Kit and the Ion OneTouch 2 system (ThermoFisher). Template-positive ion sphere particles (ISPs) were enriched using the Ion Torrent OneTouchES. Enriched ISPs were loaded on a PGM 314R v2 chip and sequenced using the Ion PGM Hi-Q View Sequencing Reagents (ThermoFisher). Raw sequencing data were processed using the Torrent Suite software v5.0 (ThermoFisher), as well as the coverage analysis and variant caller (VC) v5.0 plugins. Processed reads were aligned to the human reference genome (hg38) (60). All identified variants and the depth of coverage were visually confirmed by the Integrative Genomic viewer (IGV, Broad Institute, MIT, Cambridge, MA). Nine % of reads were filtered out by post processing. The filtered sequences, average read length 176 bp, were converted to fastq format and analyzed for non-homologous end joining (NHEJ)-mediated mutations using the CRISPResso pipeline (24, 61). Sequence reads were compared to the reference PCR sequence of the wild type huPGRN gene, GenBank M75161.1 (Fig. 1C-E).

### Cellular proliferation

Proliferation of H69 and ΔhuPGRN-H69 cells in response to exposure to 100 nM recombinant *Ov*-GRN-1 (r*Ov*-GRN-1) (17, 21) was quantified using the impedance-based xCELLigence real time cell analysis (RTCA) approach (ACEA Biosciences, San Diego, CA). The r*Ov*-GRN-1 peptide (∼10 kDa) was concentrated using Centripep with cut-off 3 kDa (Eppendorf) and resuspended in low salt solution, Opti-MEM. The absorbance at 205 nm and concentration of r*Ov-*GRN-1 was determined by using a Nanodrop 2000c spectrophotometer (ThermoFisher) (62). Five thousand cells/well were seeded in 16 well-E-plates (ACEA) in H69 complete media. E-plates was inserted in the xCELLigence DP platform at 37°C, 5% CO_2_ and changes in impedance reflecting cell adhesion and proliferation record at intervals of 20 min for 24 hrs. On the following day, the medium was removed and replaced with H69 complete medium supplemented with r*Ov*-GRN-1 at 100 nM. Cellular proliferation was monitored with data for the wild type H69, r*Ov*-GRN-treated H69, and ΔhuPGRN-H69 with and without r*Ov*-GRN-1treatment displayed as change of impedance (Cell Index) for 48 hours, normalized to wild type H69 (reference cell line in the assay) (RTCA Software 1.2, ACEA) (63).

### Isolation and characterization of exosomes

The H69 or ΔhuPGRN-H69 cells were treated with 100 nM r*Ov*-GRN-1 in H69 complete media with 10% exosome depleted-FBS. Forty-eight hours later, were harvested the supernatant from cell culture of H69 with or without, and ΔhuPGRN-H69 with or without liver fluke granulin. The supernatants were collected at 48 h following addition of r*Ov*-GRN-1 and cellular debris pelleted and removed by centrifugation 2,000×*g*, 15 min. The supernatant was filtered through a 0.22 μm pore size membrane (Millipore, Billerica, MA), mixed with 0.5 volume of tissue culture total exosome isolation reagent (Invitrogen, catalog no. 4478359), and incubated for 16 h at 4°C. Thereafter, the exosome pellet (isolated by centrifugation at 10,000×*g*, 4°C, 60 min) was re-suspended in ×PBS (64). Exosomes were investigated in co-culture assays and the exosomal RNA (exRNA) and protein (exProtein) of the exosomes were investigated following extraction of exosomal contents using the Total Exosome RNA and Protein Isolation kit (ThermoFisher).

The identification of protein markers on the isolated exosomes was undertaken using the commercially available Exo-Check Exosome Antibody Array kit (System Biosciences, Palo Alto, CA): the array included 12 pre-printed spots and was used with the cognate antibodies specific for the exosome markers CD63, CD81, ALIX, FLOT1, ICAM1, EpCam, ANXA5 and TSG101. The array included the following controls: a GM130 cis-Golgi marker to monitor cellular contamination, a positive control spot derived from human serum exosomes, and a blank spot as a background control. The membrane was developed with SuperSignal West Femto Maximum Sensitivity Substrate (ThermoFisher Scientific) and visualized and quantified on the ChemiDoc imager (Bio-Rad) (65) (66). The intensity of exosome protein markers were read (gray scale) (ChemiDoc), and subtracted with negative background intensity. The signal intensity for each protein was plotted and compared (Fig. 3B). The exProtein was also confirmed for exosome marker expression by western blot analysis against anti-CD9 and CD81 (Abcam). Briefly, 10 µg of the exProtein was separated on gradient (4-12%) SDS-PAGE gel and transferred to nitrocellulose membrane (Bio-Rad). After blocking with 5% skim milk in Tris-buffered saline (TBS)-Tween for 60 min, the membrane was incubated with specific antibody against CD9 or CD81, followed (after washing) by anti-rabbit HRP-linked secondary antibody (DAKO Corporation catalog no. P0448) diluted 1 in 2,000. Signals from ECL substrate were detected by chemiluminescence (Amersham Bioscience, Uppsala, Sweden) and captured and analyzed (ChemiDoc, Bio-Rad) (Fig. 3C).

For immunofluorescence staining of exosome particles from H69 cells were proceeded by culturing the cells in glass slide chamber overnight before fixed at 4°C in ice-cold methanol for 10 min, washed 3 times in 1× phosphate-buffered saline (PBS), and permeabilized in 0.1% Triton X-100/PBS for 10 min at room temperature. Nonspecific binding was blocked with 0.5% Tween-20/PBS containing 1% bovine serum albumin (BSA) for 30 min. The primary antibodies against fluoroflore-488 labelled-CD81 was incubated for 60 min at room temperature. The incubated cells were washed 3 times in 1×PBS. Before visualization by confocal microscope, we stained the cell nucleus with DAPI. The exosome particles were observed and particle sized were measures in H69 cytoplasm (Fig. 3A)

### Quantitative real time PCR

Total RNA and exRNA either from H69 or ΔhuPGRN-H69 cells were isolated using RNAzol (Molecular Research Center, Inc.) or total exosome RNA isolation kit (ThermoFisher) following the manufacturer’s instructions. One microgram of RNA was treated for DNase, then used for reverse transcription by an iScript cDNA synthesis kit (Thermo Fisher Scientific). Real time PCR was performed in ABI7300 Real time PCR machine using the SsoAdvanced Universal SYBR Green Supermix (Bio-Rad). The PCR reaction consisted of 5 µl SsoAdvance SYBR Green PCR master mix, 0.5 µl of 10 µM forward and reverse primers, and 2 µl of 5 times diluted template cDNA in a total volume of 10 µl. The thermal cycle was initiation cycle at 95°C for 30 sec followed by 40 cycles of annealing at 55°C for 1 min. Samples were analyzed in at least 3 biological replicates (various cell passages) and in typical reactions. The human glyceraldehyde-3-phosphate dehydrogenase (GAPDH) transcript was run parallel with human granulin (huPGRN) and used for gene normalization. The differential granulin transcript fold change was calculated by formula 2^-ddCt^ (67). The specific primers for huPGRN and GAPDH are as follows: PGRN-F: 5’-atgataaccagacctgctgcc-3’, PGRN-R: 5’-aaacacttggtacccctgcg-3’, GAPDH-F; 5’-tgtagttgaggtcaatgaaggg-3’ and GAPDH-F 5’-tgtagttgaggtcaatgaaggg-3’. The fold change was obtained by comparing treated-samples with the untreated control (indicated as a value of 1, 100%). The means and standard deviations of differential transcript expression were calculated by independent Student’s *t-*tests using GraphPad Prism software (La Jolla, CA).

### Western blot and densitometry

Protein lysates or exosomal protein (exProtein) from H69 and ΔhuPGRN-H69 cells were prepared using M-PER mammalian protein extraction reagent (ThermoFisher) and exosome protein isolation kit (ThermoFisher, catalog 4478545), respectively, following the manufacturer’s protocols. Protein concentrations were determined using the Bradford assay (68). Ten micrograms of cell lysate or 20 µg of exProtein was separated on gradient SDS-polyacrylamide gel (4-12% Bis-Tris, Invitrogen) and transferred to nitrocellulose membrane (Trans-Blot Turbo, Bio-Rad). Signals were identified with a chemiluminescence substrate (GE Healthcare) following probing with antibodies specific for huPGRN (Abcam, catalog ab 108608) or GAPDH (Sigma-Aldrich, catalog G9545) and HRP conjugated-secondary antibodies. Expression levels of huPGRN were imaged using the FluroChem system (Bio-techne, Minneapolis, MN) and relative expression levels of huPGRN investigated after GAPDH level normalization. Differences compared using independent Student’s *t-*tests.

### Cholangiocarcinoma (CCA) gene expression panel

A predesigned Cholangiocarcinoma Pathway Panel (88 targets) (PrimePCR™, Bio-Rad, Hercules, CA, catalog no. 100-2531) was used to investigate the CCA gene expression profile of transcripts in the cargo of exosomes released H69 cells exposed to liver fluke granulin. One microgram of total RNA from exosome samples was converted to cDNA (Supermix iScript kit, Bio-Rad). A one in 10 dilution of cDNA was used for qPCRs with a final concentration of 1*×* SsoAdvanced universal SYBR super mix (Bio-Rad) and 1*×*PrimePCR assays for the designated target. Reactions were performed (three technical replicates) at 10 μl final volume, using the iQ5 real time PCR system (Bio-Rad) starting with activation at 95°C for 2 min, followed by 40 cycles of denaturation at 95°C, 5 s, and annealing/elongation at 60°C, 30 s. Specificity of target amplification was confirmed by melting-curve analysis. Controls for evaluating reverse transcription, RNA quality, genomic DNA contamination and kit performance were included on the array (Bio-Rad). The reference genes GAPDH, HPRT1 and TBP were assayed for relative gene expression to normalize for variation in quantity of input mRNA. The DNA template served as a positive real-time PCR control for the corresponding gene assay. PrimePCR Analysis Software (Bio-Rad) was used for analysis of differential fold changes of target genes.

### Uptake of PKH26-labeled exosomes by cholangiocytes

To investigate whether r*Ov*-GRN-1-treated H69 use exosomes to communicate CCA-conducive mRNAs to adjacent naïve cells, exosomes were labeled with PKH26 (Fluorescent cell linker kit, Sigma-Aldrich) (69, 70). In brief, isolated exosomes (6.25 µg) from culture media were re-suspended in one ml of diluent C, to maintain cell viability with maximum dye solubility and staining efficiency (71) and labeled by addition of PKH26 to 2 µM final concentration. The samples were mixed gently and incubated for 5 min (periodic mixing), after which 2 ml of 1% BSA was added to bind excess dye. The mixture containing PKH26-labeled exosomes was subjected to precipitation (Tissue culture exosome isolation reagent, ThermoFisher), as above, after which the exosomes were suspended in H69 complete medium and co-cultured with H69 or ΔhuPGRN-H69 cells at 80-90% confluence in wells of poly-*L*-lysine-coated microscope slide chambers (Lab-Tek II). A negative control group was co-cultured with exosomes not stained with PKH26. After incubation with PKH26-labeled exosomes for 90 min at 37°C in 5% CO_2_, the cells were stained with NucBlue Live Cell Stain ReadyProbes (Invitrogen) and visualized by confocal fluorescence microscopy (Zeiss Cell Observer SD Spinning Disk Confocal Microscope, Carl Zeiss Microscopy, Thornwood, NY). A negative control for inhibition of endocytosis was included, which involved addition of Pitstop 2, an inhibitor of clathrin-mediated endocytosis (Abcam, catalog ab120687) (32) at 30 μM to recipient cells for 15 min, before co-culture with labeled-exosomes (Fig. 5A-C).

### Human MAPK phosphorylation array

To investigate whether CCA-associated functional RNAs in exosomes were translated by naïve H69 cells (both wild type and the ΔhuPGRN), the cells were incubated with the exosomes as described above. For this, 5×10^5^ cells were seeded into a well of 24-well plate and maintained overnight in H69 complete medium (72). Thereafter, exosome particles were dispensed into each well and cultured for 24 h. The cells were harvested with cell scraper, then washed 3 times with 1*×* PBS, before resuspended in Lysis buffer provided in the Human MAPK Phosphorylation kit). The cell lysate was measure for protein concentration by Bradford reagent (Bio-Rad) prior to proceed to Human MAPK Phosphorylation array (Abcam) following the manufacturer’s instructions. In brief, protein sample was incubated with each array overnight at 4°C on an orbital shaker. The unbound proteins were removed, and the arrays washed 3X with washing buffer. Arrays were incubated with the horse radish peroxidase anti-rabbit IgG antibody for 120 min at room temperature. Thereafter, arrays were washed 3X times with washing buffer prior to incubation with freshly prepared detection solution for 2 min at room temperature. The protein spots were visualized using the chemiluminescence signal (ChemiDoc Imaging System, Bio-Rad). The intensity score of each duplicated array spot was measured, and the normalized array data were calculated according to the manufacturer’s recommendation. To normalize array data, one array was defined arbitrarily as the ‘reference array’, to which the other arrays were compared. The following algorithm was used to calculate and determine the signal expression between like analytes: X(Ny) = X(y) * P1/P(y), where P1 = mean signal density of positive control spots on reference array; P(y) = mean signal density of positive control spots on array ‘y’; X(y) = mean signal density of spot ‘X’ on array for sample ‘y’; and X(Ny) = normalized signal intensity for spot ‘X’ on Array ‘y’. We used exosome derived-ΔhuPGRN-H69 cells (not exposed to r*Ov*-GRN-1) as the control to contrast with the *in vivo* translation of CCA-associated mRNAs in the treatment groups.

### Biological and technical replicates, statistics

Biological replicates represented parallel measurements of biologically discrete samples in order to capture random biological variation. Technical replicates were undertaken as well; these represented repeated measurements of the same sample undertaken as independent measurements of the random noise associated with the investigator, equipment or protocol. Responses of knockout versus wild type H69 cell line genotypes including before and after exposure to liver fluke granulin were compared using one-way ANOVA, including for multiple comparisons at successive time points. Thereafter, Tukey’s honest significance and Student’s *t* tests were performed to compare treatment groups. Cell index (CI) assays were performed in triplicate. CI was automatically registered by the RTCA software (ACEA Biosciences Inc., San Diego, CA). Means ± SD are indicated, plotted using GraphPad Prism 8 (GraphPad Software, San Diego, CA). Values of *P* < 0.05 were considered to be statistically significant.

Supplementary Figure 1 provides a schematic overview of methods and findings.

### Database accessions

Nucleotide sequence reads reported here have been assigned SRA accessions SRR9735031-34 at GenBank, NCBI, and the study has been assigned BioProject ID PRJNA556235 and BioSample accession SAMN12344265.

## Acknowledgements

We thank Drs. Griffin P. Rodgers, NIDDK, National Institutes of Health (NIH) and Yang Liu, Institute of Human Virology, School of Medicine, University of Maryland for support with exosome studies and Ion Torrent-based deep sequencing. PA was supported by the Ph.D. program at the Faculty of Medicine, Khon Kaen University and the Thailand Research Fund through the Royal Golden Jubilee Ph.D. Program, award number PHD/0111/2557 (PA, TL). We acknowledge support from award R01CA164719 (TL, AL, PJB) from the National Cancer Institute (NCI), NIH. The content is solely the responsibility of the authors and does not necessarily represent the official views of the Thailand Research Fund, the NCI or the NIH.

## Supporting information

**Supplementary Figure 1.**
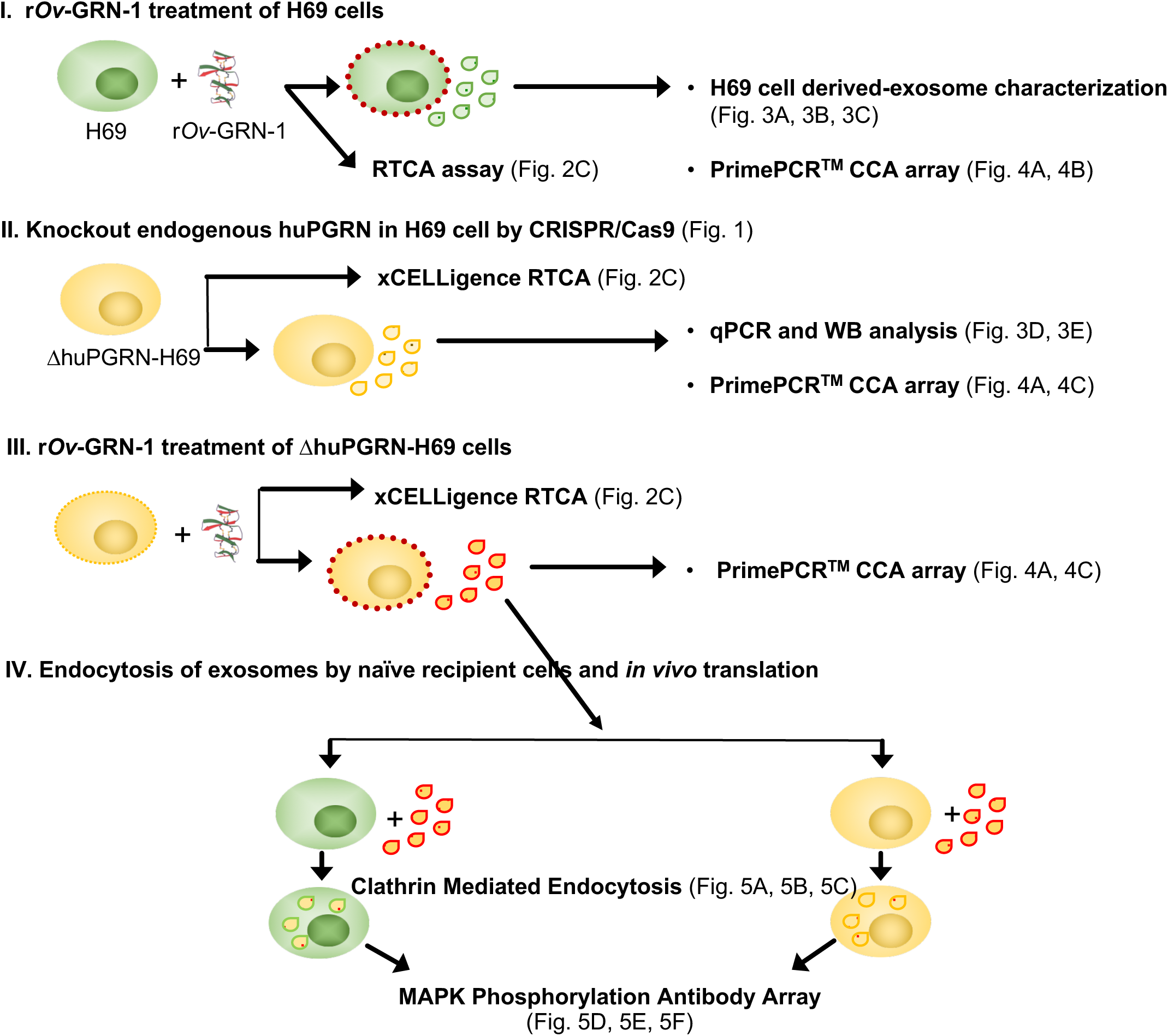
Schematic representation of experimental approach and methods, involving the use of intact and gene-edited cholangiocytes to investigate activation by liver fluke granulin and paracrine signaling by exosomes

